# Partial lesions of hypocretin neurons of the rat lateral hypothalamus affect wake and sleep regulation under increased sleep pressure

**DOI:** 10.64898/2026.10.19.759695

**Authors:** Maria Lebedeva, David Levčík, Jan Kudláček, Kristýna Malenínská, Iveta Vojtěchová, Helena Buchtová, Hana Brožka, Dominika Radostová, Erika Petránová, Přemysl Jiruška, Jan Svoboda, Štěpán Kubík, Eduard Kelemen, Aleš Stuchlík

## Abstract

Hypocretin (orexin)-producing neurons in the lateral hypothalamic area are important for sleep and wake regulation in mammals and their damage in humans manifests clinically as narcolepsy. While in previous studies experimentally induced massive loss of these neurons revealed their physiological importance, small lesions and their manifestations can uncover more subtle features of sleep regulation and help understand the delicate processes in early stages of pathogenesis of narcolepsy. We studied the effects of partial lesions (approximately 30%) of hypocretin-producing neurons in the lateral hypothalamus on sleep patterns and EEG in rats. The lesions did not change the normal sleep and wake pattern in undisturbed rats. The effects of lesions manifested under increased sleep pressure after six hours of sleep deprivation, when they led to accelerated dynamics of compensatory slow wave sleep. The lesions also led to a slight decrease in theta frequency in wakefulness and in REM sleep, which we observed during undisturbed sleep and wake behavior as well as after sleep deprivation.

**Highlights:** Partial lesions to 30% of hypocretin-producing neurons in the lateral hypothalamus did not cause detectable change to the normal undisturbed sleep and wake pattern in rats.

Under increased sleep pressure after six hours of sleep deprivation, the effects of the lesions manifested by earlier occurrence of compensatory slow wave sleep.

In the EEG of the lesioned animals, a slight shift to lower theta frequencies was observed during wakefulness and REM sleep.

## Introduction

Narcolepsy is a rare neurological disease with prevalence of 0.02-0.067% (1,2) that manifests primarily by excessive daytime sleepiness, and may be accompanied by a heterogeneous collection of other symptoms with variable severity. The presence or absence of cataplexy, i.e. sudden loss of muscle tone without loss of consciousness, divides the disease into two diagnostic units – narcolepsy type 1 (NT1), when cataplexy is present, and narcolepsy type 2 (NT2), when cataplexy is absent. NT2 often develops into NT1 and can be seen as an early stage of the pathology. Other symptoms of narcolepsy include rapid-eye-movement sleep (REMS) intrusions into wakefulness, sleep onset REMS periods (SOREMPs) (3–5), distortions of night’s sleep by hypnagogic hallucinations, sleep paralysis, repeated awakenings during night, increased time in stage N1, and decreased time in stage N3 of non-REM sleep (6). The underlying cause of narcolepsy is damage to the hypocretin-producing neurons in the lateral hypothalamic area (LHA) (7) triggered by an autoimmune reaction (8–12). Hypocretins (orexin-A and orexin-B) are excitatory neuropeptides acting via G-protein coupled hypocretin receptors 1 (Hcrtr1) and hypocretin receptors 2 (Hcrtr2). Hcrtr2 has similar affinity to both peptides and localizes to areas involved in arousal such as tuberomammilary nucleus and dorsal raphe (13). Selective Hcrtr2 agonists show promise in clinical trials for narcolepsy treatment (14,15).

The research in laboratory animals has typically focused on modelling the most severe manifestations of narcolepsy as seen in NT1. Lesions (16), as well as genetic manipulations (17–19) were used to massively damage the hypocretin-producing system. These manipulations led to a set of symptoms, which included dysregulation of sleep and wake cycles and short latency of REMS similar to human SOREMPs (16–20). The genetic models also reproduced episodes of sudden inactivity, muscle atonia, and postural collapse resembling cataplexy (17–19). However, mice with genetic ablation of hypocretin-producing neurons also displayed late-onset obesity despite lower food intake (18), suggesting that the role of hypocretin-producing neurons is not limited to sleep-wake regulation, but extends to metabolic, autonomous, and affective functions as well (13). To understand finer details of functions of the hypocretin-producing neurons in sleep regulation and to investigate early etiology of narcolepsy requires more subtle manipulations of the hypocretin system than massive lesions.

Here we set out to characterize sleep and wake behavior in rats after partial lesions of approximately one third of hypocretin-producing neurons in the LHA. In lesioned and sham-operated animals, we characterized sleep and wake patterns during 24 hours of undisturbed behavior, and during recovery from six hours of sleep deprivation. We also characterized electroencephalography (EEG) during wakefulness and in different sleep stages. We observed no significant effects of lesions on sleep structure and duration during spontaneous behavior. In conditions of increased sleep pressure after the sleep deprivation, we observed enhancement and altered dynamics of compensatory slow wave sleep in lesioned animals compared to the sham-operated controls. In the lesioned animals, we also observed decreased frequency in theta range in EEG during wakefulness and REM sleep.

## Materials and Methods

Nineteen adult male rats of Long-Evans strain (2-4 months old) obtained from the breeding colony of the Institute of Physiology of the Czech Academy of Sciences were used in the experiment. Animals were divided into a lesioned group (N = 9) and a sham-operated group (N = 10). The rats were housed in standard plastic boxes with free access to food and water in a controlled light and dark cycle with lights on at 7 a.m. and lights off at 7 p.m. The temperature was maintained at 22±2 °C and humidity at 50±20%. All animal procedures were approved by the Committee for the Ethical Treatment of Animals and Animal Welfare at the Institute of Physiology, Czech Academy of Sciences (Project No. 80/2016) and complied with the Animal Protection Act of the Czech Republic and European Union directive 2010/63/EU.

Electrophysiological methods for sleep and wake monitoring followed previously used procedures (21,22). Bone screws were used as electrodes for EEG recording and were attached to a Mill-Max connector through insulated nickel-chromium wires (75 µm diameter, California Fine Wire Company, Grover Beach, CA, USA). A wire with a free end stripped of insulation served as an electromyography (EMG) electrode. Prior to lesions and electrode implantation, the rats were anesthetized by isoflurane and placed into a stereotaxic frame (Stoelting, Wood Dale, IL, USA). During the surgery, the rats were kept under general anesthesia (1.5 – 2.5% isoflurane). The scalp was removed, the skull was exposed, and openings for injections were drilled in the skull. For the rats in the lesioned group, lesions were produced by bilateral injections of hypocretin-2-saporin (Hcrt2-SAP) conjugate (30 ng per hemisphere, Advanced Targeting Systems, Escondido, CA, USA) stereotaxically aimed at the LHA (3.3 mm anterior, ±1.7 mm lateral relative to bregma, and 8.2 mm ventral relative to dura; (20)). Rats of the sham-operated group received injections of sterile 0.9 % saline solution into the LHA. Injections were made using a Hamilton syringe at a flow rate of 0.075 µl/min in a total volume of 0.5 µl of solution per hemisphere. After injections, five holes were drilled into the skull for the electrode screws. The screws in the frontal area (3 mm anterior, ±2 mm lateral to bregma) and parietal area (4 mm posterior, 2 mm lateral to bregma) were used for EEG recording. Two screws in the occipital bone were used as grounding and reference electrodes. The EMG electrode was inserted under the skin into the dorsal neck region. The Mill-Max connector was fixed to the skull with dental cement (Dentalon, Kulzer GmbH, Hanau, Germany). The wound was sutured and treated with povidone-iodine (Betadine 2%, EGIS, Prague, Czechia) and antibiotic ointment (Framykoin, Zentiva Group, Prague, Czechia) to prevent inflammation. The animals were left to recover until the wound was fully healed.

Twelve to fourteen days after intracerebral injections and electrode implantation, EEG and EMG signals were recorded, and sleep and wake patterns were characterized in each rat in a 48-hour-long recording session. This experimental protocol followed previously used methodology (23). The recording was performed in a familiar recording box (40 cm x 30 cm, with 60 cm walls) with food and water available *ad libitum*. The first 24 hours represented a normal undisturbed sleep and wake cycle, with lights on at 7:00 (7:00 a.m.) and lights off at 19:00 (7:00 p.m.). Next, the animals were sleep deprived for six hours by a “gentle handling” procedure during the first half of the light period (from 7:00 to 13:00). The recovery from sleep deprivation was characterized during the following 18 hours (13:00 to 7:00 the following day; Fig. 1A).

**Figure 1.**
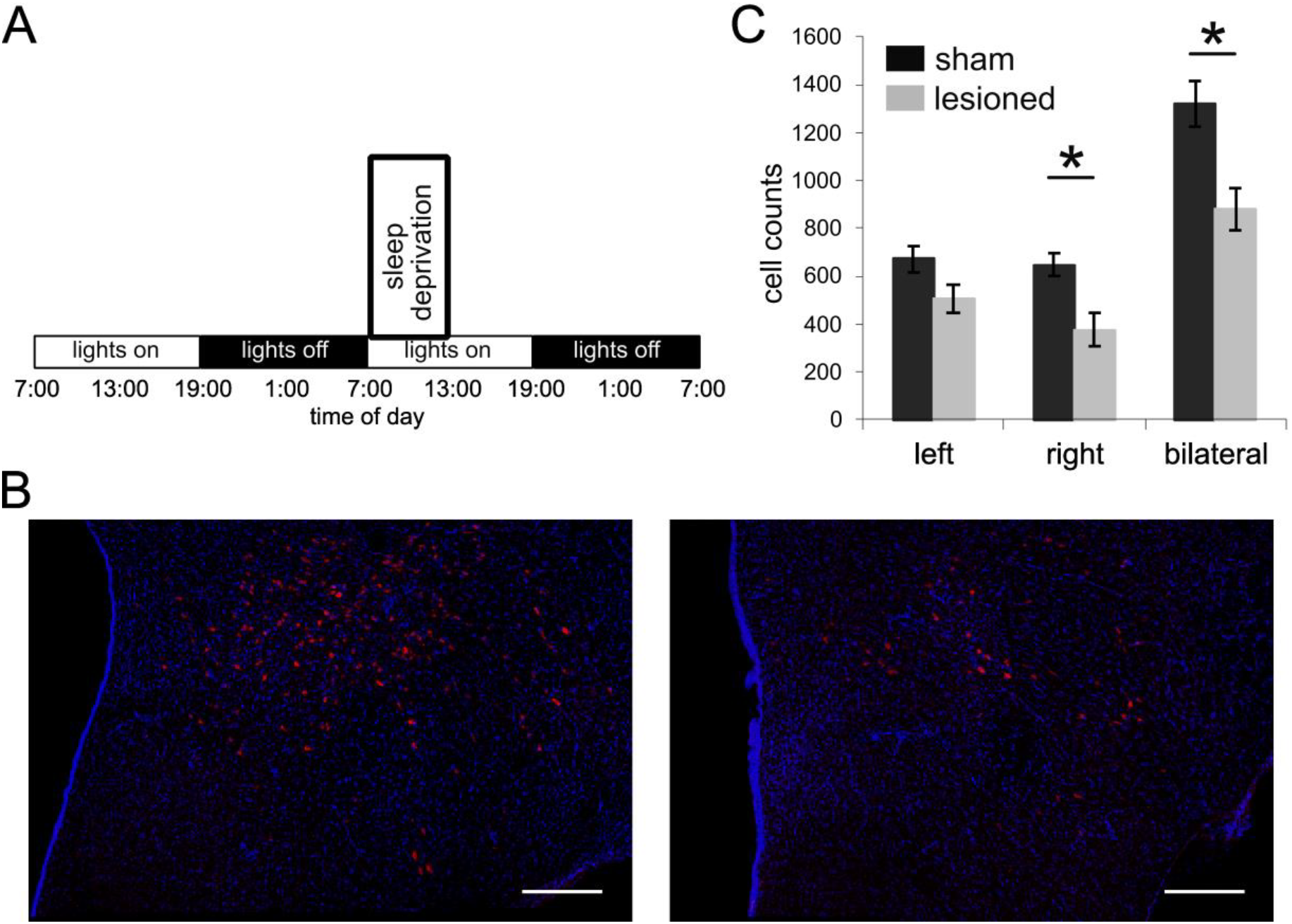
A) Experimental protocol. After lesions of LHA or sham procedure and following 12 – 14 days of recovery period, EEG and EMG activity was recorded continuously during 48 hours in a recording cage. The lights were on from 7:00 to 19:00, lights were off from 19:00 to 7:00 the next day. For the first 24 hours, the animals were undisturbed and natural sleep and wake pattern was characterized. The second day started with six hours of sleep deprivation (7:00 – 13:00). The sleep and wake pattern during the following 18-hour period of compensatory sleep was characterized. B) Example of histological staining of LHA in sham-operated rat (left panel) and lesioned rat (right panel). Hypocretin antibodies are shown in red, cell nuclei are marked by DAPI and shown in blue. The white scale bars mark 300 µm. C) Cell counts of neurons in LHA stained by hypocretin antibodies. (Data are presented as mean ± SEM. * p < 0.01)

At the time of recording, the rats were connected to a recording system consisting of Intan RHD2000 controller and RHD2132 headstage amplifiers (Intan Technologies, Los Angeles, CA, USA). EEG signals were filtered between 1 and 300 Hz and recorded at 2000 samples per second. EMG signals were filtered between 30 and 300 Hz and also sampled at 2000 samples per second. Recordings were evaluated offline by experimenters blind to the lesion conditions, who used a custom-made Matlab program (MathWorks, Natick, MA, USA). Each 10-second interval was manually scored as wake, slow wave sleep (SWS), REMS, or preREMS, based on EEG and EMG signals using standard criteria (24) as in our previous work (21,22). Briefly, intervals with high activity on the EMG electrode, often accompanied by theta activity in EEG were scored as wake intervals. Intervals with low EMG activity and high proportion of slow frequencies below 4.0 Hz in EEG were scored as SWS. Intervals with low EMG activity and high theta EEG activity were scored as REMS. Intervals with intermediate features between SWS and REMS were scored as preREMS. For each scored interval, the Fast Fourier Transform (FFT) of EEG was performed, and the power spectrum was calculated and stored for further analysis.

After the completion of the experiments, the rats were anesthetized by a ketamine and xylazine mix (50 mg/kg) and perfused with 0.2 M phosphate buffered saline (PBS), followed by 4% paraformaldehyde (PFA) in PBS. Their brains were further fixed in PFA for 24 hours and underwent cryoprotection in 10%, 20%, and 30% sucrose in PBS. The brains were then frozen on dry ice and stored at -80 °C. Next, the brains were sectioned at 40 µm with a Leica 1850 cryostat and stored in a cryoprotective solution. Every fourth section was used for lesion assessment. For hypocretin neuron detection, sections were washed with PBS and incubated in a blocking buffer with 0.3% Triton X-100 and 10% bovine serum in PBS for an hour. Sections were then treated with a primary hypocretin antibody (1:500, Anti-Orexin A, Abcam ab6214) for 48 hours at 4 °C. After washing with PBS, sections were incubated with anti-rabbit antibodies conjugated with Alexa Fluor Plus 555 (1:1000, goat anti-rabbit IgG, Invitrogen A32732), counterstained with DAPI (1:20000), mounted, and sealed with nail polish. For analysis, every fourth section of the lateral hypothalamus was examined, producing 7-11 confocal stacks per rat. Images were captured using a Leica TCS SP8 microscope with a 20× immersion objective, comprising 26 layers per stack. The Alexa555 signal was excited at 554 nm and filtered through a 565-665 nm bandpass. Neuron counts were analyzed using ImageJ software (25) on the maximal Z-stack projection.

For statistical assessment, SPSS Statistics package (IBM, Armonk, NY, USA) was used. Paired t-tests and two-way ANOVAs with Newman-Keuls test for post hoc comparison were used. Significance was accepted at p < 0.05.

## Results

The histologically determined number of hypocretin-containing neurons in the lesioned group (880.6±85.9) was significantly lower than in the sham-operated group (1320.3±94.9; t(17) = 3.407; p = 0.003; Fig. 1B and 1C). The mean reduction in number of neurons in the lesioned group was 33.3%.

The sleep/wake pattern and sleep architecture were first characterized during 24 hours of undisturbed spontaneous behavior, with lights on during the first 12 hours (light phase) and lights off during the second 12 hours (dark phase). Lesioned and sham-operated rats did not differ in distribution of time spent in different wake and sleep stages during the 24 hours (Fig. 2, panels A-D). The part of the day (light phase vs. dark phase) had a significant effect on the proportion of time spent awake and in different stages of sleep. No significant effects of the lesion or interaction were found. Specifically, the proportion of time spent in the wake state was lower during the light phase compared to the dark phase for both lesioned and sham-operated animals. A two-way ANOVA indicated no interaction (F(1, 34) = 0.202; p = 0.656), a significant effect of the part of the day (F(1, 34) = 152.943; p < 0.001), and no significant effect of lesion (F(1, 34) = 0.006; p = 0.938). The proportion of time spent in each one of different sleep stages (SWS, REMS, and preREMS) was higher in the light phase compared to the dark phase. For SWS, there was no interaction (F(1, 34) = 0.1; p = 0.754), a significant effect of the part of the day (F(1, 34) = 88.647; p < 0.001), and no effect of lesion (F(1, 34) = 0.067; p = 0.798). For REMS, there was no interaction (F(1, 34) = 0.126; p = 0.725), the effect of the part of the day was significant (F(1, 34) = 43.123; p < 0.001), and there was no significant effect of the lesion (F(1, 34) = 0.036; p = 0.852). For preREMS, again, the effect of interaction was not significant (F(1, 34) = 0.048; p = 0.827), there was a significant effect of the part of the day (F(1, 34) = 14.111; p = 0.001), and there was no significant effects of lesion (F(1, 34) = 1.109; p = 0.301).

**Figure 2.**
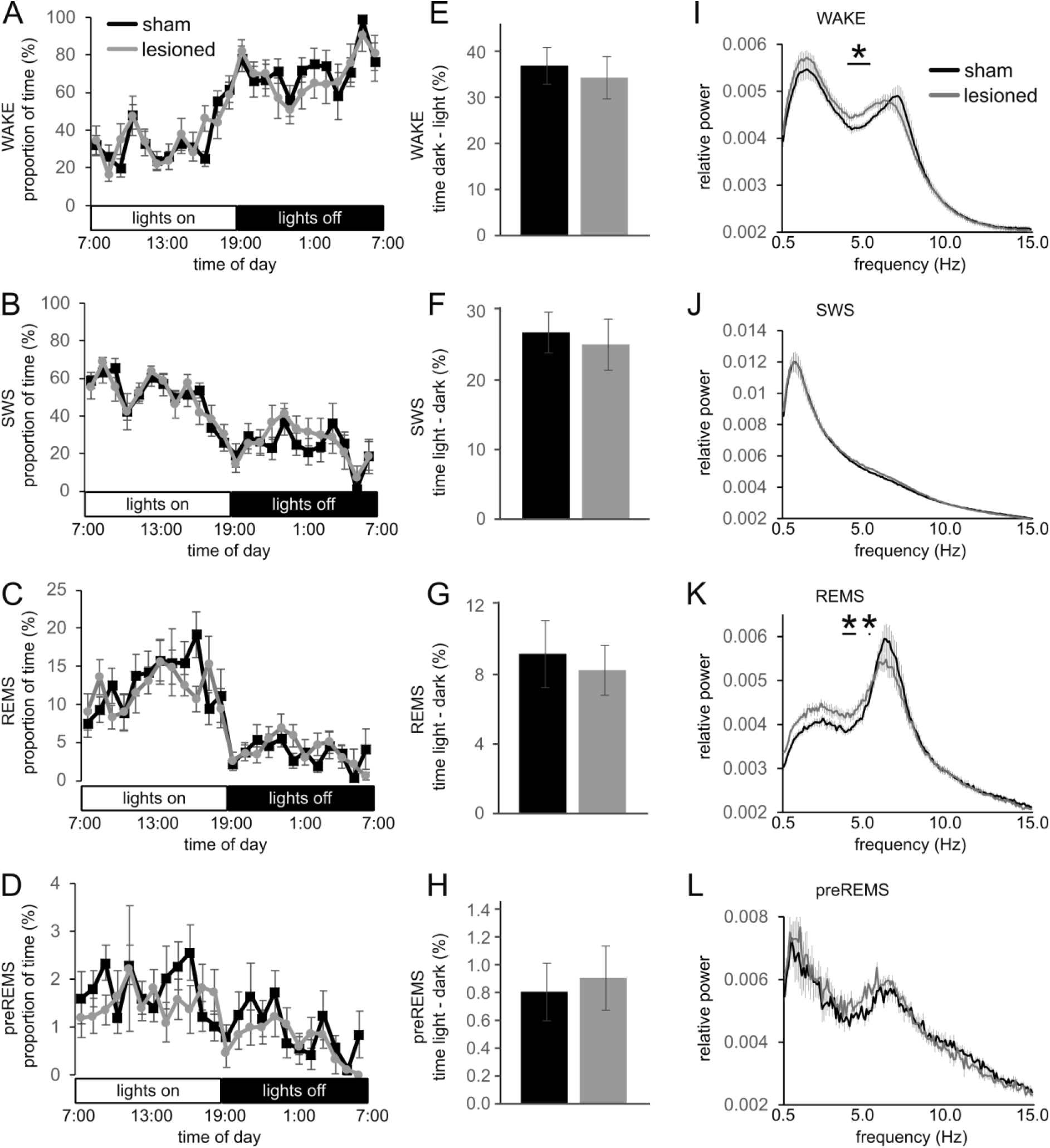
Sleep and wake pattern during 24 hours of spontaneous undisturbed behavior. A–D) Distribution of time spent in different wake and sleep stages for one-hour intervals. A) wake, B) SWS, C) REMS, D) preREMS. E–H) Difference between the light phase and dark phase in time spent awake (E), in SWS (F), in REMS (G) and in preREMS (H). I–L) Power spectra for EEG during different wake and sleep stages. I) wake, J) SWS, K) REMS, L) preREMS. (Data are presented as mean ± SEM. In panels I and K horizontal lines and asterisks mark range of frequencies with significant difference in power between the two experimental groups. * p < 0.05)

Previous studies reported that differences in sleep pattern between the light and dark phase are reduced after extensive lesions of hypocretin receptor-expressing cells in the LHA (16). We therefore calculated the difference between the light and dark phase in time spent awake (Fig. 2E). The lesioned and sham-operated groups did not significantly differ in this parameter (t-test, p > 0.05). Similarly, lesions had no significant effect on the difference between the light and dark phase in SWS time (Fig. 2F), REMS time (Fig. 2G), and preREMS time (Fig. 2H) (t-tests, p’s > 0.05).

The frequency composition of EEG signals in different wake and sleep stages during 24 hours of undisturbed spontaneous behavior was characterized using the FFT analysis and power spectra (Fig. 2I-L). During the wake state, we observed a tendency for a slight decrease in theta frequencies in the lesioned rats compared to sham-operated rats (Fig. 2I). This tendency was manifested by a significant increase of theta power in lower theta range (4.3, 4.5-5.6 Hz) in the lesion animals (p’s < 0.05). During SWS, there was no significant difference in power spectra between the two groups (Fig. 2J). In REMS, we observed in lesioned animals increased power at lower theta and delta frequencies. The difference was significant for frequencies 4.0, 4.1, 4.3-4.7, 5.5 Hz (p’s < 0.05, Fig. 2K). The number of detected preREMS episodes was too small for reliable statistical analysis (Fig. 2L).

The power of slow oscillations, which is related to sleep pressure, was analyzed during SWS periods. In undisturbed spontaneous behavior, the power of slow oscillations in SWS decreased during the light phase, that is the rats’ inactive period, for both sham-operated and lesioned groups (Fig. 4A). For quantitative analysis, the average SWS slow oscillation power during the first hour and the last hour of the light phase was analyzed (Fig. 4B). Two-way ANOVA revealed no interaction between time and treatment (F(1, 30) = 0.134; p = 0.717). There was a significant effect of time (F(1, 30) = 6.637; p < 0.05), and no significant effect of lesion (F(1, 30) = 0.115; p = 0.737).

Next, we studied the effect of the lesions on compensatory sleep following six hours of sleep deprivation (SD). The sleep pattern during each hour after the SD was compared to the corresponding hour before the SD (i.e. time interval 24 hours earlier). Cumulative histograms allow comparison of lesioned and sham-operated groups and show the cumulative amount of increased time of compensatory sleep and decreased time of wakefulness after SD (Fig. 3A-D). Compensatory sleep was observed in both groups of animals. Both groups spent less time awake (Fig. 3A) and more time in SWS and REMS (Fig. 3B,C) after SD compared to pre-SD levels (t-tests, p’s < 0.05). Neither sham-operated nor lesioned rats significantly increased their preREMS duration after the SD (Fig. 3D). The dynamics of temporal structure of changes in wake time and sleep time after SD differed between the two groups of rats. Thus, changes in sleep pattern after the SD were studied in 3-hour intervals (Fig. 3E-H). The difference between the lesioned and sham-operated groups was strongest during the first 3-hour interval of the dark phase, i.e. at the beginning of the rats’ active part of the day. During this interval, the lesioned animals significantly reduced the time spent in the wake state (t(17) = 2.603; p = 0.019; Fig. 3E), while the time in the SWS was significantly increased (t(17) = - 2.992; p = 0.008; Fig. 3F) compared to the sham-operated group. Increases of time spent in REMS and preREMS after the beginning of the dark phase were also observed in both groups of rats, but the differences between lesioned and sham-operated animals were not significant. Next, we compared proportions of time spent in different wake and sleep stages between the light and dark phases of the day (Fig. 3M-P). Lesioned rats had significantly smaller differences between the light and dark phases compared to the controls in REMS (t(17) = 2.237; p = 0.039; Fig. 3O) and preREM sleep (t(17) = 2.247; p = 0.038; Fig. 3P).

**Figure 3.**
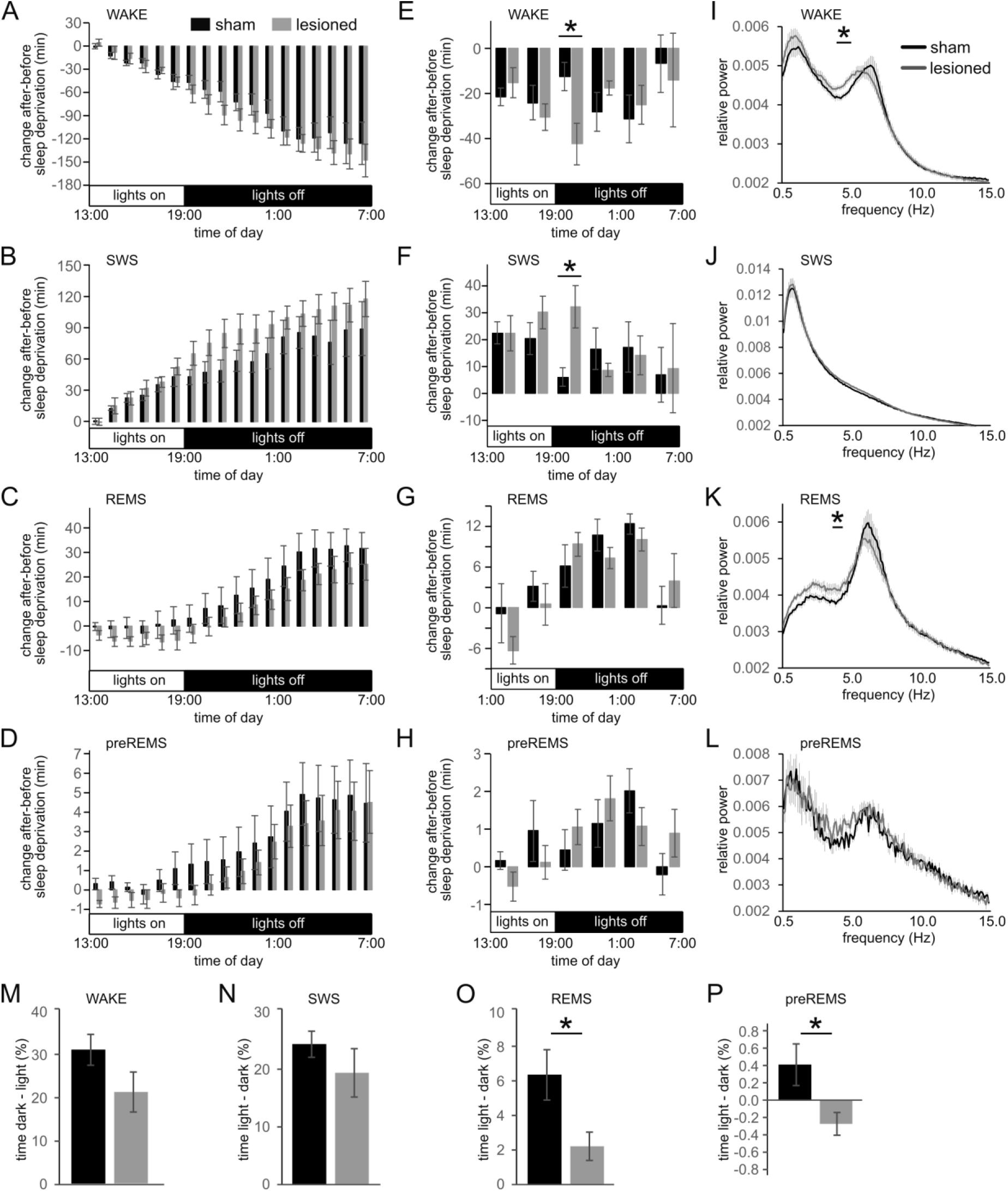
Compensatory response to six hours of sleep deprivation. A–D) Cumulative change in time spent in different wake and sleep stages after sleep deprivation in comparison to the same one-hour time intervals 24 hours earlier. A) wake, B) SWS, C) REMS, D) preREMS. E–H) Change in time spent in different wake and sleep stages after sleep deprivation in comparison to the same intervals 24 hours earlier is shown for 3-hour intervals. E) wake, F) SWS, G) REMS, H) preREMS. I–L) Power spectra for EEG during different wake and sleep stages. I) wake, J) SWS, K) REMS, L) preREMS. M–P) Difference between the light phase and dark phase in time spent awake (M), in SWS (N), in REMS (O) and in preREMS (P). (Data are presented as mean ± SEM. In panels I and K horizontal lines and asterisks mark range of frequencies with significant difference in power. * p < 0.05)

The frequency composition of EEG in different wake and sleep stages after SD (Fig. 3I-L) displayed a similar pattern as during undisturbed sleep and wake behavior. During the wake state, there was a tendency for a slight decrease in theta frequencies in the lesioned group (Fig. 3I). This tendency was manifested by a significant increase of power in lower theta range (4.6, 4.8-5.4 Hz) in the lesioned group (p’s < 0.05). During SWS, there was no significant difference in power spectra between the two groups (Fig. 3J). In REMS, we observed that lesions led to lower relative power at the theta peak accompanied by increased power at lower theta and delta frequencies. The difference was significant for frequencies 4.0, 4.2, 4.4-4.7 Hz (p’s < 0.05, Fig. 3K). Statistical comparison was not performed for the small number of detected preREMS episodes (Fig. 3L).

After six hours of SD, the power of slow oscillations in SWS decreased during the light phase for both the sham-operated and lesioned groups (Fig. 4A). The average SWS slow oscillation power during the first hour after sleep deprivation and the last hour of the light phase was analyzed (Fig. 4C). Two-way ANOVA revealed no interaction between time and treatment (F(1, 33) = 0.263; p = 0.612). There was significant effect of time (F(1, 33) = 6.145; p < 0.05), and no significant effect of lesion (F(1, 33) = 2.370; p = 0.133). To assess the effect of SD on slow oscillations, the power of slow oscillations during the first hour of the light phase in undisturbed conditions was compared to the first hour after SD. There were no significant differences in slow-oscillation power after SD in neither control (t(17) = - 0.584; p = 0.567) nor lesioned animals (t(15) = -0.137; p = 0.893).

**Figure 4.**
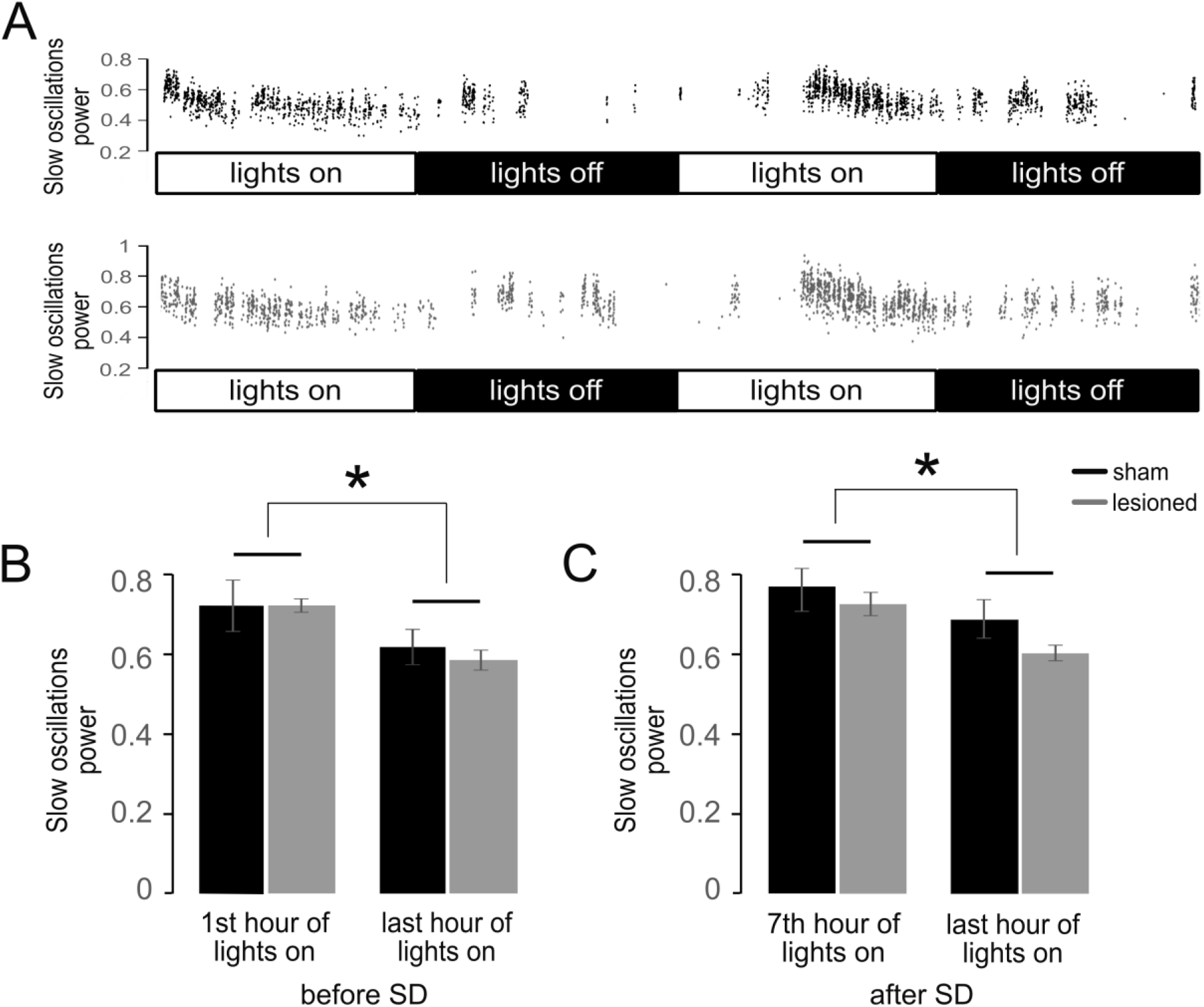
Changes in the power of slow oscillations during SWS. A) Two examples of dynamic changes in the power of slow oscillations in SWS during 24 hours of undisturbed sleep and wake, followed by six hours of sleep deprivation and 18 hours of recovery. The upper panel shows an example from a sham operated rat, and the lower panel represents an example of lesioned rat. B, C) Quantitative analysis of changes of slow oscillation power between beginning and end of light phase before (B) and after (C) sleep deprivation. (Data are presented as mean ± SEM. * p < 0.05)

Sleep onset REM sleep periods (SOREMPs) are a characteristic feature of narcolepsy in people. To assess the potential presence of a similar phenomenon in our model, the latency of REMS onset (duration of wake-REMS intervals) was studied. The distributions of the duration of wake-REMS intervals did not significantly differ between the sham-operated and lesioned animals. In laboratory animals, SOREMPs were previously identified as periods when REMS occurred within less than two minutes after a wake period that lasted for at least two minutes (16). The overall number of SOREMPs did not significantly differ between the lesioned group and the sham-operated group (t(17) = -0.572; p = 0.575) and the numbers of detected SOREMPs were very low in both groups (data not shown).

## Discussion

We induced partial, relatively small lesions of the lateral hypothalamus of rats to model pathology associated with narcolepsy. These lesions destroyed approximately one third of hypocretin-producing neurons and did not lead to detectable changes in sleep architecture during 24 hours of spontaneous sleep and wake behavior. However, the effect of the lesions manifested after six hours of sleep deprivation. Under these conditions of increased sleep pressure, the lesioned animals compensated for lost sleep faster than the sham-operated rats. At the onset of the dark phase, the lesioned rats spent less time awake and more time in slow wave sleep compared to sham-operated animals. Examination of EEG showed a slight decrease in theta frequencies during wakefulness and REMS, which manifested both before and after sleep deprivation.

We induced and histologically confirmed partial lesions of hypocretin-producing neurons in LHA by injection of 30 ng of Hcrt2-SAP per hemisphere. Effects of Hcrt2-SAP lesions on neuronal tissue in the lateral hypothalamus have been previously thoroughly characterized (16,20,26). In these studies, Hcrt2-SAP was applied in concentrations substantially higher than in our work (90ng/0.5µl and 490ng/0.5µl) and the effects of the lesions on different types of neurons in the posterior hypothalamus, which includes lateral hypothalamus and adjacent nuclei, have been described. As expected, the lesions decreased the number of hypocretin-immunoreactive neurons (by 88% at Hcrt2-SAP concentration of 90 ng/0.5 µl, and by 91% at Hcrt2-SAP concentration of 490 ng/0.5 µl), but the effects of the lesions were not limited to hypocretin-immunoreactive neurons. A similar loss of melanin-concentrating hormone (MCH)-containing neurons was observed, as well as a loss of ADA-immunoreactive neurons in the tuberomammillary nucleus (16). On the other hand, α-melanocyte stimulating hormone (α-MSH)-producing neurons were spared (16). Based on these data, we used much lower dose of Hcrt2-SAP to minimize damage to neuronal types other than hypocretin-producing neurons and prevent off-target effects such as disrupted feeding. The relevance of hypocretin-producing neurons to narcolepsy is obvious, as these neurons are lost and hypocretin levels decreased in narcoleptic individuals and ablation of these neurons recapitulate the symptoms in animal models (18,19). Furthermore, selective Hcrtr2 agonists relieve the narcoleptic symptoms in clinical trials (14,15). MCH cells are not damaged in human narcoleptics (7) and MCH knock out mice do not show signs of narcolepsy (27), although dual ablation of hypocretin and MCH neurons increased wakefulness and caused more severe cataplexy and sleep attacks compared to ablation of hypocretin neurons alone (28). Functional role in metabolism, rather than sleep regulation, was suggested for these cells (26). Histaminergic (ADA-immunoreactive) neurons in tuberomammillary nucleus have been implicated in arousal, as they activate in wake and discharge much less in sleep (29–31). Their damage by Hcrt2-SAP may therefore affect sleep and wake behavior. Although we cannot rule out involvement of other neuronal types, given the low dose used and demonstrated reduction of hypocretin-producing neurons in LHA, we argue that the loss of hypocretin-producing neurons is the most probable cause of the effects observed in this study.

NT1 and NT2 in humans differ in the extent of damage to hypocretin neurons in the lateral hypothalamus. NT1 is associated with a massive, 85-95% loss of hypocretin-producing LHA neurons (7), close to undetectable levels of hypocretin (Hcrt) mRNA in the hypothalamus, undetectable levels of Hcrt-1 and Hcrt-2 peptides in the brains of patients (32), and undetectable levels of Hctr-1 in cerebrospinal fluid (33). NT2, on the other hand, is characterized by less severe, approximately 33% loss of LHA hypocretin neurons (34) and Hcrt-1 levels of approximately 50 pg/ml in cerebrospinal fluid, which is higher compared to NT1, but substantially lower than approximately 210 pg/ml in healthy individuals (33). Cases of NT2 may represent early stages of NT1 with less pronounced symptoms (2,35). In people with low hypocretin in cerebrospinal fluid, about one third may develop cataplexy, and thus NT1, years later. This makes NT2 a particularly interesting subject of study, useful for understanding early etiology and subtle primary pathological mechanisms of the disease. The extent of the lesions in our study corresponds to cell loss seen in NT2.

The role of hypocretin and hypocretinergic neurons in sleep regulation has been broadly studied in laboratory animals. Experimental approaches included lesions of hypocretinergic neurons in lateral hypothalamus (16,20), hypocretin gene knockout (17), the study of naturally occurring hypocretin receptor mutations (36), and genetic ablation of hypocretin-producing neurons (18,19). The lesion studies were typically associated with massive destruction of hypocretinergic neurons. Lesions of more than 60% of hypocretinergic neurons in LHA in rats led to substantial changes in sleep architecture: a decrease in wake time and an increase in amount of sleep during the dark (active) part of the rat day (16,20). Similarly, the amount of time in REMS and SWS increased and the amount of time awake decreased in the dark phase of a day in hypocretin knockout mice (17). Interestingly, in human patients there are reports of increase of daytime REMS, but no changes in overall amount of sleep and no increase in daytime SWS (37,38). In one study, the amount of SWS was higher in untreated patients with narcolepsy than in controls (39). The hallmark of human NT1 is cataplexy, which, in animal models, was possibly observed in the form of “narcolepsy episodes” in hypocretin knockout mice (17), but not in LHA lesions by Hcrt2-SAP (16). SOREMPs – another characteristic symptom of human narcolepsy – were observed in both hypocretin knockout mice (17) and in rats after LHA lesions by Hcrt2-SAP (16). Finally, both mice (18) and rats (19) with genetic ablation of hypocretin-producing neurons show signs of cataplexy, reduced wakefulness and SOREMPs. Nonetheless, unlike the monogenic animal models (17,36), human narcolepsy is typically not familial and may be caused by environmental factors acting on a susceptible genetic background (40).

Experiments with smaller lesions of hypocretin-producing cells in LHA in laboratory animals are scarce. In rats with lesions of < 30% of hypocretin-producing neurons, the amount of time spent in wake and sleep stages and the number of SOREMPs were not different from controls (16). This is very similar to our observations. However, we observed changes in sleep architecture after six hours of sleep deprivation, where lesioned animals compensated more quickly for the lost sleep with more SWS and less wakefulness compared to control animals. This manifested especially during the first three hours after the onset of darkness, which is the beginning of the active part of the day for the rats, when most narcolepsy symptoms are observed in humans and animal models. Our findings are consistent with observations in humans that after 24 hours of sleep deprivation, NT1 patients tend to compensate with more SWS during recovery period compared to control subjects, while the amount of compensatory REMS is not different from controls and time in stage N2 of non-REM sleep is decreased in patients (41,42).

We observed a decrease in the peak of theta (5-10 Hz) in power spectra and a decrease in theta frequencies in wakefulness, both before and after sleep deprivation. A similar tendency was observed in REMS. Mukai et al. (43) observed decrease of theta frequency power (4.0-5.0 Hz) during REM sleep in narcoleptic patients compared to controls, but a similar effect was not reported in a different human study by (41). Cholinergic activity in the medial septum and diagonal band of Broca plays an important role in theta rhythm generation. Medial septal neurons receive strong innervation from hypocretin neurons, contain abundance of hypocretin receptors 2, and septo-hippocampal neurons are excited by hypocretin (44,45). Thus, attenuation of hypocretin input caused by lesions in our study might indirectly interfere with generation of theta rhythm. Hypocretinergic neurons affect theta rhythm also through modulation of the dopaminergic system. Combined knockout of Hcrtr1 and Hcrtr2 on dopaminergic neurons led to suppression of theta power (46), consistent with our findings. Interestingly, the effect of hypocretin was dependent on hypocretin receptor type, and knockout of Hcrtr2 alone led to increase in theta power (46).

In addition to changes in theta power, human narcoleptic patients displayed less “fast delta” activity (1.5-3.4 Hz) and more “slow delta” activity (0.5-1.0 Hz) in REM sleep (43). In non-REM sleep, narcoleptics displayed an increase in sigma frequencies (11.0-11.5 Hz), which overlap with sleep spindle activity, and a small increase in delta power (1.0-1.5 Hz). Similar increases in delta and sigma frequencies were observed in narcoleptics after sleep deprivation (41). Such changes were not observed in our study.

In summary, we aimed to characterize the early physiological changes associated with partial loss of hypocretin-producing neurons. After lesions of approximately 30% of these neurons, we observed only subtle changes in sleep pattern, and the hallmark changes accompanying massive loss of hypocretin neurons in narcolepsy – SOREMPs and cataplexy – were not observed. These observations suggest robust organization, sturdiness, and redundancy of the hypocretin system. This redundancy may hamper the early detection of developing pathology in clinical settings. We also identified that the changes associated with mild damage manifested at the time of increased sleep pressure after sleep deprivation. These changes may represent early signs of deteriorating sleep-wake regulation in NT2. They may provide screening methods at an early stage, before critical symptoms manifest, when protective cure may prevent further damage and development of the full syndrome.

## Acknowledgements

Work was supported by the Czech Science Foundation grant number 17-08596S, awarded to Jan Korábečný and Štěpán Kubík, and ERDF-Project Brain Dynamics, No. CZ.02.01.01/00/22_008/0004643. We are grateful to Dr. Jitka Bušková and Dr. Tomáš Petrásek for useful comments on earlier versions of the manuscript and to Alexa Yang for proofreading English text.

